# Targeting homologous recombination repair to potentiate fluoroquinolone efficacy in *Mycobacterium abscessus*

**DOI:** 10.1101/2025.10.24.684385

**Authors:** Shuai Wang, Md Shah Alam, Buhari Yusuf, Jingran Zhang, Ziwen Lu, Xiaofan Zhang, H.M. Adnan Hameed, Mst. Sumaia Khatun, Chunyu Li, Lijie Li, Aleksey A. Vatlin, Aweke Mulu Belachew, Xirong Tian, Yamin Gao, Cuiting Fang, Rogers Peng Zhang, Jun Li, Xinyue Wang, Liqiang Feng, Li Wan, Tianyu Zhang

**Author notes:** Correspondence: Tianyu Zhang. These authors contributed equally to this work (Shuai Wang, Md Shah Alam). (S.W.); (J.Z.); (H.M.A.H); (Y.G.); (M.S.A.); (B.Y.); (Z.L.); (X.Z.); (M.S.K.); (C.L.); (A.M.B.); (L.F.); (L.W.); (L.J.); (A.A.V.); (X.T.); (C.F.); (R.P.Z.); (X.W.); (J.L.).

## Abstract

Synthetic antibiotics, fluoroquinolones (FQs), are extensively important treatment options for fast growing mycobacterial infections. However, their widespread use has led to decreased efficacy due to intrinsic or acquired resistance. FQs exhibit antimicrobial activity by stabilizing type II topoisomerases-generated DNA breaks, thereby inducing lethal double-strand breaks (DSBs) and leading to bacterial death. Escalating resistance among mycobacterial pathogens highlights the need to elucidate resistance mechanisms and identify novel strategies to restore or enhance the drug activity. Here, we revealed homologous recombination (HR) as a key determinant of FQ tolerance in *Mycobacterium abscessus*, a notoriously drug-resistant pathogen. Through a transposon mutagenesis screen, we identified that disruption of *adnB* - an HR-associated gene, markedly sensitized *M. abscessus* to FQs. This finding prompted a systematic dissection of DSB repair pathways in *M. abscessus*. Targeted deletion of HR core components (*adnB, recO, recA, recR, ruvB*) significantly increased FQ susceptibility *in vitro*, while inactivation of alternative DSB repair pathways - single-strand annealing (SSA) and non-homologous end joining (NHEJ) - had no effect, revealing a unique reliance on HR for FQ tolerance. Furthermore, the complementation of gene knock-out *M. abscessus* strains ΔadnB, ΔrecO, ΔrecA, ΔrecR or ΔruvB with the corresponding genes from *M. abscessus*, as well as homologous genes from *Mycobacterium tuberculosis* restored the resistance phenotype, indicating a conserved HR-dependent tolerance mechanism in *M. abscessuss*. In a murine infection model, genetic abrogation of HR significantly improved the therapeutic efficacy of FQs against *M. abscessus*, demonstrating translational relevance. Collectively, our findings position HR as a conserved and actionable vulnerability in *M. abscessus*, and provide a compelling rationale for developing HR-targeted adjuvant therapies to resensitize drug-refractory strains to FQ treatment.

**Impact statement:** Fluoroquinolone (FQ) resistance poses a major challenge in the treatment of *Mycobacterium abscessus* infections. This study identifies homologous recombination (HR) as a key defense mechanism that limits FQ efficacy across multiple mycobacterial species. Genetic inhibition of HR not only sensitizes *M. abscessus* to FQs *in vitro* but also enhances antibiotic activity *in vivo*. These results highlight HR repair as a promising therapeutic target to potentiate FQ activity and combat drug resistance in *M. abscessus* infections.

## Introduction

The rapidly growing *Mycobacterium abscessus* complex has emerged as a major healthcare-associated opportunistic pathogen responsible for diverse infections^1^. Recent epidemiological reports indicate increasing prevalence of *M. abscessus* in East Asia^2, 3^, with infections now comprising approximately 17.2% of clinical non-tuberculous mycobacteria (NTM) isolates worldwide ^4^. Among rapidly growing NTMs, *M. abscessus* is the most pathogenic and drug-resistant species, frequently causing chronic pulmonary infections, particularly in immunocompromised individuals, as well as skin and soft tissue infections ^5, 6^. The clinical presentation of *M. abscessus* lung disease often resembles pulmonary tuberculosis (TB), complicating diagnosis and management ^5^. Rising incidence and intrinsic multidrug resistance make *M. abscessus* infections a formidable therapeutic challenge, necessitating the development of novel therapeutic strategies and the identification of new drug targets to enhance treatment efficacy ^7^.

Fluoroquinolones (FQs) remain highly effective antimicrobial agents for the treatment of drug-resistant TB and are increasingly employed against NTM infections, particularly in macrolide-resistant *M. abscessus* strains ^8, 9^. Moxifloxacin (MFX), a fourth-generation FQ, plays a pivotal role in non-tuberculosis mycobacteria regimens treatment, exhibited relative good *in vitro* activity against *M. abscessus* clinical isolates in previous studies ^10^. The antibacterial activity of FQs is mediated through inhibition of type II topoisomerases, specifically DNA gyrase and topoisomerase IV, which are important for DNA replication, transcription, and chromosome segregation ^11^. Mycobacteria possess DNA gyrase as their sole type II topoisomerase, uniquely enabling the introduction of negative DNA supercoils. FQs stabilize the covalent DNA–enzyme cleavage complex formed by type IIA topoisomerases, thereby preventing re-ligation of cleaved DNA strands and causing an accumulation of double-strand breaks (DSBs), ultimately resulting in bacterial cell death ^12^.

Efficient repair of DSBs is crucial for maintaining genomic stability in mycobacteria. The repair of mycobacterial DSB is mediated by three distinct routes: (i) homologous recombination (HR), a high-fidelity RecA-dependent mechanism that restores sequence integrity; (ii) non-homologous end joining (NHEJ), an error-prone Ku–LigD–dependent process that ligates broken DNA ends without homology; and (iii) single-strand annealing (SSA), a RecA-independent pathway requiring RecBCD and strand-annealing proteins such as RecO, which often results in deletions between homologous repeats ^13, 14^.

Transposon mutagenesis has proven highly effective in dissecting the genetic basis of bacterial drug resistance and identifying potential therapeutic targets. This method involves random insertion of transposons into the bacterial genome, generating a diverse mutant library that enables high-throughput screening to identify genes critical for antibiotic susceptibility, bacterial growth, and stress tolerance. Such insights are instrumental in development of novel antimicrobial strategies. For instance, in *M. tuberculosis*, Gupta *et al*. discovered that transposon insertion disrupting the nonclassical transpeptidase LdtMt2 led to increased susceptibility of *M. tuberculosis* to the β-lactam antibiotic amoxicillin ^15^. Follow-up studies demonstrated that meropenem effectively inhibits both Ldt_Mt2_ and an additional transpeptidase ^16^, providing the rationale for combining meropenem with clavulanate, a regimen that has shown efficacy against drug-resistant *M. tuberculosis* ^17^.

In this study, we constructed a transposon mutant library of *M. abscessus* to identify genes associated with altered susceptibility to FQs. Our results demonstrate that disruption of the HR repair pathway significantly increases the susceptibility of *M. abscessus* to FQs, suggesting the HR repair pathway as a potential therapeutic target. This research paves the way for developing novel combinatorial therapeutic strategies to address the challenges posed by this persistent mycobacterial pathogen and offers new opportunities for more effective treatment of most naturally resistant *M. abscessus* infections.

## Results

### Disruption of *adnB* sensitizes *M. abscessus* to FQs

To investigate genetic determinants of intrinsic FQs resistance in *M. abscessus*, we performed a transposon mutagenesis screen of >5,000 mutants (Figure 1A). One mutant, designated S27, exhibited markedly increased sensitivity to FQs (Figure 1B and Table 1). The minimum inhibitory concentrations (MICs) of two FQs, levofloxacin (LEV) and MFX, against strain S27 were reduced to 1/16 of those observed for wild-type *M. abscessus* (Table 1). Sequencing revealed a transposon insertion within *MAB_3515c*, located 337 bp downstream of the start codon (Figure 1C). Sequence analysis showed that MAB_3515 shares 59.21% amino acid identity with *M. tuberculosis* AdnB, a DNA helicase involved in HR, suggesting that *MAB_3515c* is the homolog of *adnB* (Figure S1).

**Table 1.** MICs of various drugs against different mycobacterium strains.

| Strains<sup>a</sup> | HR repair process | Drugs<sup>b</sup>/MICs<sup>c</sup> (µg/mL) |  |  |  |
| --- | --- | --- | --- | --- | --- |
|  |  | LEV | MFX | MMC | BLM |
| Mab | / | 16 | 8 | 4 | 1 |
| S27 | / | 1 | 0.5 | 1 | 0.5 |
| ΔadnB | End resection helicase-nuclease | 1 | 0.5 | 1 | 0.5 |
| ΔadnBCMab |  | 8 | 4 | 4 | 1 |
| ΔadnBCMtb |  | 4 | 2 | 2 | 1 |
| ΔrecO | Resecting the ends to form single-stranded DNA | 1 | 0.25 | 1 | 1 |
| ΔrecOCMab |  | 16 | 8 | 4 | 1 |
| ΔrecOCMtb |  | 16 | 8 | 2 | 1 |
| ΔrecA | Facilitating strand invasion | 1 | 0.25 | 1 | 0.5 |
| ΔrecACMab |  | 16 | 8 | 4 | 1 |
| ΔrecACMtb |  | 16 | 8 | 2 | 1 |
| ΔrecR | Essential for HR-mediator activity for RecA loading | 1 | 0.5 | 1 | 0.5 |
| ΔrecRCMab |  | 16 | 8 | 4 | 1 |
| ΔrecRCMtb |  | 16 | 8 | 2 | 1 |
| ΔruvB | Resolving the resulting Holliday junctions to restore genomic integrity | 1 | 0.25 | 1 | 0.5 |
| ΔruvBCMab |  | 16 | 8 | 4 | 1 |
| ΔruvBCMtb |  | 16 | 8 | 2 | 1 |

<sup>a</sup> Mab, wild-type *M. abscessus*; S27, Mab strain with transposon (Tn) insertion in *adnB*; ΔadnB, *adnB* deletion strain; ΔrecO, *recO* deletion strain, and so on; ΔadnBCMab, complemented strain expressing Mab*adnB*; ΔadnBCMtb, complemented strain expressing Mtb*adnB*, and so on. <sup>b</sup> LEV, levofloxacin; MFX, moxifloxacin; MMC, mitomycin; BLM, Bleomycin. <sup>c</sup> MIC is defined as the lowest concentration of a drug that inhibits visible bacterial growth.

**Figure 1:**
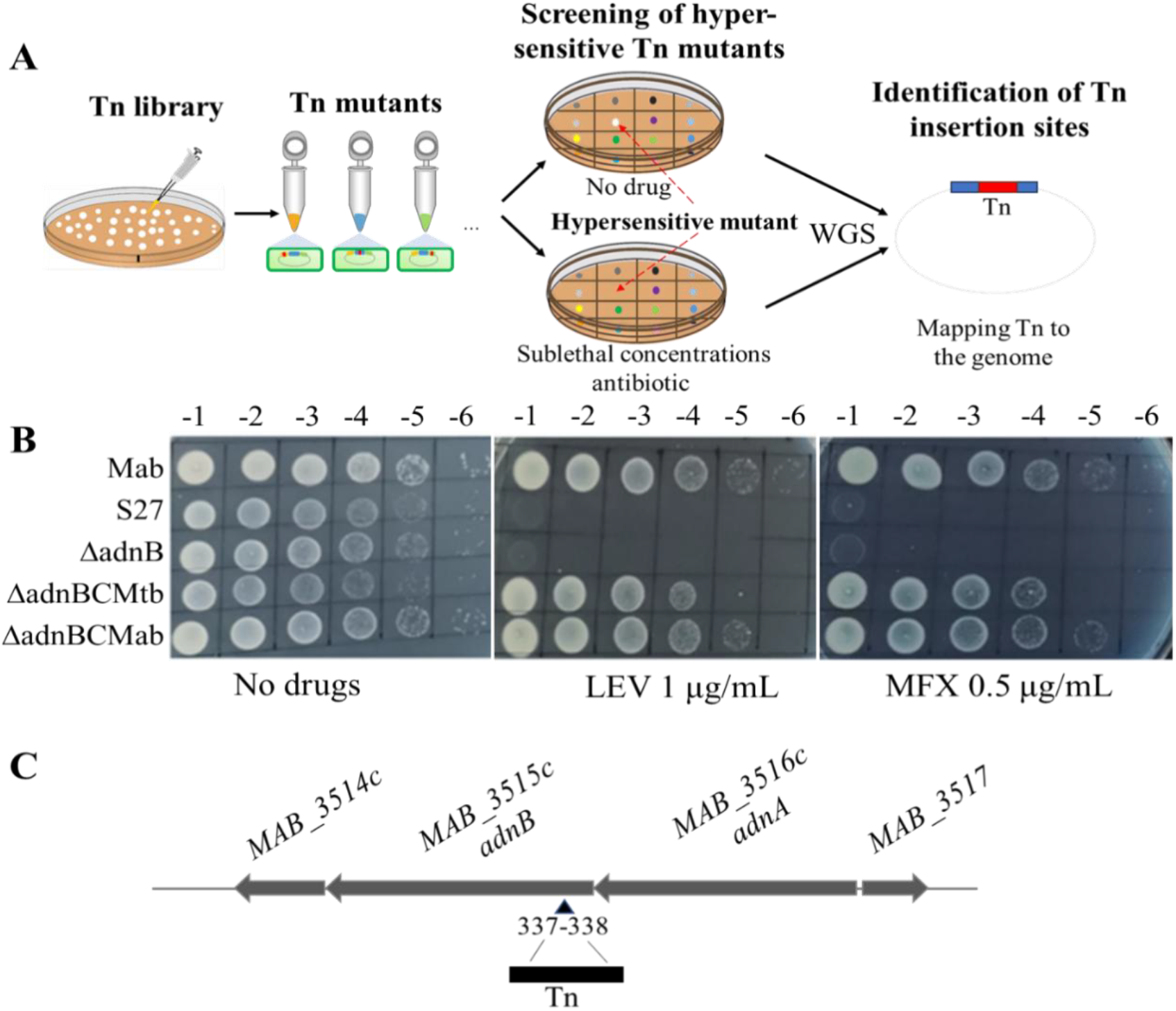
Disruption of the *adnB* results in increased sensitivity of *M. abscessus* to FQs. (A) Schematic overview of the transposon mutagenesis screening strategy used to identify genes involved in intrinsic resistance to FQs. (B) Susceptibility of different *M. abscessus* strains to FQs. (C) Diagram showing the transposon insertion site within the *adnB* (*MAB_3515c*) gene. The insertion occurs at a ‘TA’ dinucleotide, 337 bp downstream of the start codon. Tn, transposon; WGS, whole-genome sequencing. Mab, wild type *M. abscessus*; S27, Transposon (Tn) insertion site containing mutant; ΔadnB, *adnB* deletion strain; ΔadnBCMab, complemented strain expressing Mab*adnB*; ΔadnBCMtb, complemented strain expressing Mtb*adnB*. LEV, levofloxacin; MFX, moxifloxacin. Representative data from three independent experiments are shown.

To confirm the phenotype and exclude polar effects of transposon insertion, we constructed an in-frame *adnB* gene knockout strain in *M. abscessus* (ΔadnB) and subsequently complemented it with the *adnB* gene derived from *M. abscessus* or *M. tuberculosis*, resulting in ΔadnBCMab and ΔadnBCMtb, respectively. Drug susceptibility testing revealed that the phenotype of ΔadnB was consistent with that of strain S27, showing hypersensitivity to LEV and MFX, while complementation with either *M. abscessus* or *M. tuberculosis adnB* gene restored the wild-type resistance phenotype (Figure 1B, Table 1). Notably, deletion of *adnB* did not alter susceptibility to antibiotics with unrelated mechanisms of action (Table S1), suggesting that *adnB* specifically modulates resistance to FQs in *M. abscessus*.

### Suppression of the HR pathway increases FQs sensitivity in *M. abscessus*

Given the AdnB’s role in HR and the increased FQs sensitivity observed upon its disruption, we hypothesized that other key components of the HR repair pathway may similarly influence FQ susceptibility in *M. abscessus*. HR repair in mycobacteria involves several coordinated steps: recognition of DSBs, 5’–3’ end resection to generate single-stranded DNA, strand invasion into a homologous template, DNA synthesis, and resolution of Holliday junctions to restore genome integrity (Figure 2A) ^14^. To assess the role of HR in mediating FQs resistance, we constructed deletion mutants for key genes involved in distinct stages of HR—*recA, recO, recR*, and *ruvB*—as well as corresponding complementation strains. All HR-deficient mutants displayed pronounced hypersensitivity to FQs, with MICs reduced to 1/16–1/32 of that in the wild-type strain (Figure 2B, Table 1). Importantly, disruption of the HR repair pathway had minimal impact on the growth of *M. abscessus* (Figure S2), indicating that the increased sensitivity to FQs is not due to a general growth defect. Complementation with either native or *M. tuberculosis* orthologs restored the resistance phenotype, demonstrating functional conservation and confirming the essential role of these HR factors in maintaining FQs resistance. These findings support the concept that targeting the HR repair pathway may serve as a promising strategy to potentiate FQs activity against *M. abscessus*.

**Figure 2:**
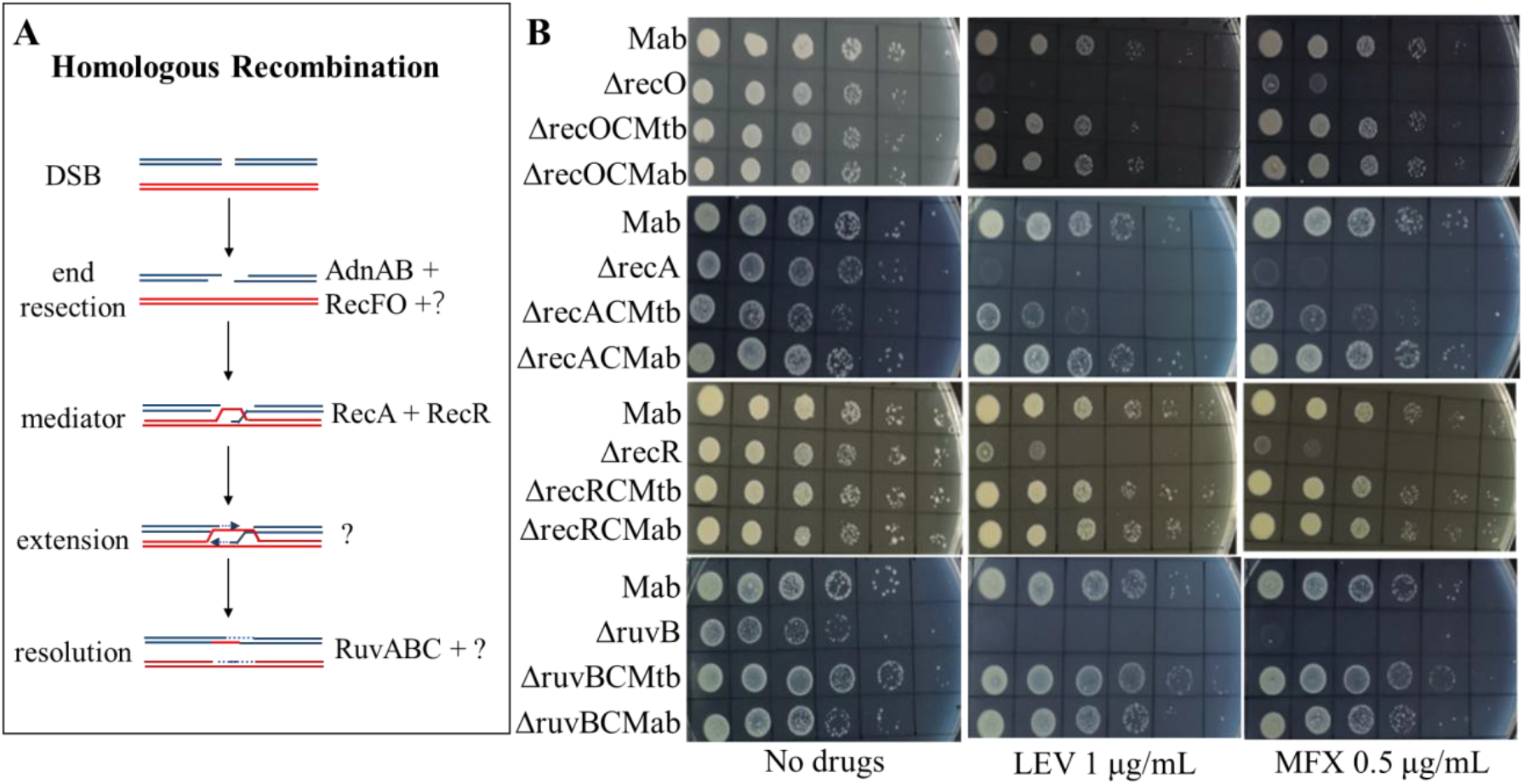
Suppression of HR enhances FQs sensitivity in mycobacteria. (A) Schematic representation of the HR-mediated DSB repair pathway in mycobacteria. (B) Drug susceptibility of different *M. abscessus* strains to FQs. Mab, wild type *M. abscessus*; ΔadnB, *adnB* deletion strain; ΔrecO, *recO* deletion strain, and so on; ΔadnBCMab, complemented strain expressing Mab*adnB*; ΔadnBCMtb, complemented strain expressing Mtb*adnB*, and so on. LEV, levofloxacin; MFX, moxifloxacin. Representative results from three independent experiments are shown.

To explore whether HR disruption generally sensitizes *M. abscessus* to different DNA-damaging agents, we tested the susceptibility of the same mutants to mitomycin C (MMC) and bleomycin (BLM). As summarized in Table 1, HR-deficient strains displayed increased sensitivity to both MMC and BLM, though to a lesser extent than with FQs, with MICs reduced by only 2-to 4-fold compared to wild-type. This differential effect may reflect the distinct nature of DNA lesions induced: while FQs predominantly generate DSBs, BLM primarily induces single-strand breaks ^18^, and MMC introduces interstrand cross-links that obstruct replication forks ^19^. Accordingly, the HR pathway plays a less prominent role in repairing damage caused by MMC and BLM compared to FQ-induced DSBs.

### Disruption of the SSA and the NHEJ repair pathway does not affect the susceptibility of *M. abscessus* to FQs

In *M. abscessus*, DSBs are repaired not only through HR but also via SSA and NHEJ ^14^. To further investigate the impact of SSA and NHEJ DSB repair pathways on the sensitivity of *M. abscessus* to FQs, we constructed *ku, ligD*, or *recC* gene knockout strains in *M. abscessus* (Δku, ΔligD, ΔrecC) and assessed their sensitivity to FQs. In contrast to HR-deficient mutants, the MICs for LEV and MFX against the *ku, ligD*, and *recC* knockout strains remained unchanged compared to the wild-type strain (16 and 8 μg/mL, respectively) (Table S2). These findings suggest that disrupting NHEJ and SSA does not affect the sensitivity of *M. abscessus* to FQs.

### HR suppression increases the *in vivo* susceptibility of *M. abscessus* to FQs

According to Clinical and Laboratory Standards Institute (CLSI) breakpoints, HR-deficient *M. abscessus* strains are considered susceptible to FQs ^20^. To validate the therapeutic potential of targeting HR repair enzymes, we evaluated the efficacy of MFX in a murine aerosol infection model using ΔadnB, ΔrecA, and ΔruvB strains (Figure 3A). As expected, MFX treatment did not significantly reduce bacterial burden in the lungs and spleens of mice infected with wild-type *M. abscessus* (Figure 3B and C). In contrast, mice infected with HR-deficient strains (ΔadnB, ΔrecA, ΔruvB) showed significant reductions in bacterial load upon MFX treatment, consistent with *in vitro* findings. Among the knockout strains, *Δ*adnB exhibited a slightly higher MIC to MFX compared to *Δ*recA and ΔruvB, which correlated with a relatively modest reduction in lung colony-forming units (CFUs) *in vivo* (Figure 3B). These data collectively indicate that disruption of the HR repair pathway sensitizes *M. abscessus* to FQs *in vivo*.

**Figure 3:**
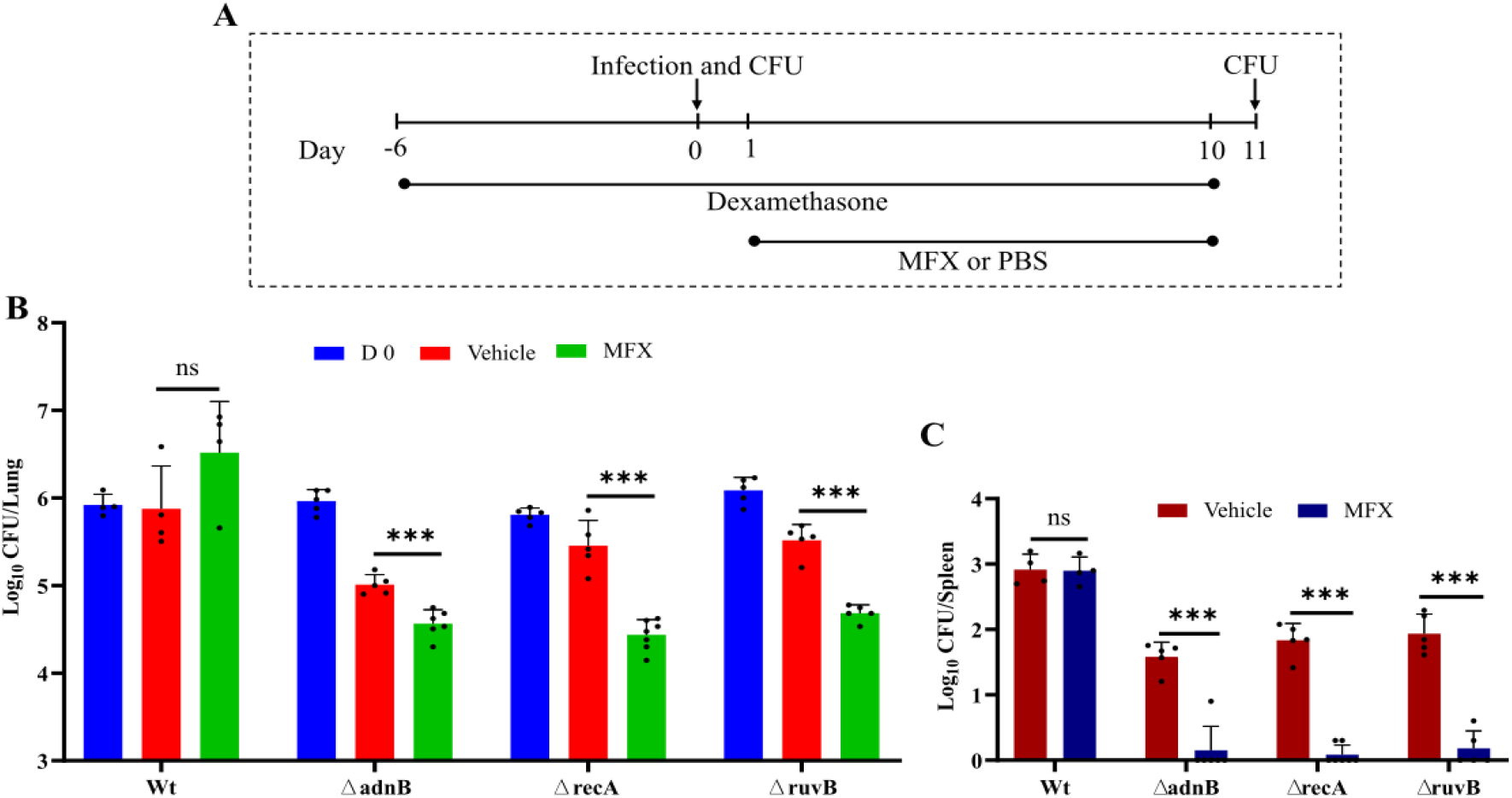
*In vivo* evaluation of *M. abscessus* susceptibility to FQ. (A) Schematic representation of the murine *M. abscessus* infection model. Mice were immunosuppressed with dexamethasone prior to infection. Mice were sacrificed at 4 hours post-infection (Day 0) to assess bacterial implantation and at 11 days post-infection (10 days post-antibiotic treatment) to evaluate the efficacy of treatment. (B) *M. abscessus* CFUs in the lungs of mice treated with moxifloxacin (MFX) versus those treated with PBS (vehicle). (C) *M. abscessus* CFUs in the spleens of mice treated with MFX versus those treated with PBS (vehicle). Each data point represents an individual mouse. Statistical significance: ns, no significant difference; ***, *P* < 0.001. Wt, wild type *M. abscessus*; ΔadnB, *adnB* deletion strain; ΔrecA, *recA* deletion strain; ΔruvB, *ruvB* deletion strain; MFX, moxifloxacin.

Interestingly, despite similar initial infection loads across all groups, mice infected with HR-deficient strains exhibited significantly reduced lung CFUs by the end of the experiment, even without MFX treatment (*P* < 0.05), suggesting impaired bacterial persistence or virulence. In contrast, bacterial loads in the lungs of untreated mice infected with wild-type *M. abscessus* remained unchanged (*P* > 0.05) (Figure 3B). A similar trend was observed in the spleen CFUs (Figure 3C), further supporting the hypothesis that disruption of the HR repair pathway compromises bacterial fitness and facilitates enhanced clearance *in vivo*. This finding underscores the dual impact of HR deficiency, both sensitizing bacteria to FQs and impairing their virulence.

## Discussion

Antibacterial synergy is a cornerstone of modern antimicrobial therapy. Rational combinations of antibiotics can significantly enhance therapeutic efficacy, limit the emergence of resistance, and potentially reduce drug-related toxicity. Understanding and applying this concept is essential for optimizing clinical treatment strategies and improving patient outcomes. A classic example is the combination of β-lactam antibiotics with β-lactamase inhibitors. β-lactam antibiotics, such as penicillin and cephalosporins, target bacterial cell wall synthesis. However, many bacteria produce β-lactamases enzymes that hydrolyze β-lactams and confer resistance. Co-administration of a β-lactamase inhibitor (e.g., clavulanic acid) protects the antibiotic from enzymatic degradation and restores its activity against resistant strains ^21^. This synergistic approach not only improves antimicrobial efficacy but also reduces the likelihood of resistance development. In this context, our study identified a novel synergistic strategy, that disruption of the HR repair pathway markedly increased the sensitivity of mycobacteria to FQs. This highlights the potential of targeting key HR-related enzymes to improve the antibacterial activity of FQs. These findings expand the understanding of FQ mechanisms of action and resistance, and offer a conceptual framework for designing synergistic therapeutic strategies, including drug repurposing and novel combination regimens for treating mycobacterial infections.

The proposed mechanism by which HR pathway disruption sensitizes mycobacteria to FQs is illustrated in Figure 4. During replication and other cellular processes, topoisomerases such as GyrA and GyrB generate transient DSBs to alleviate DNA supercoiling. FQs bind to the DNA-topoisomerase complex, forming a stabilized ternary structure that prevents re-ligation of the DNA, ultimately resulting in bacterial cell death. In response to FQ-induced DSBs, bacteria activate DNA repair mechanisms, with HR being the predominant high-fidelity repair pathway in mycobacteria. HR-mediated repair enables some bacterial cells to survive (Figure 4, bottom left). However, in the absence of functional HR, DSBs cannot be efficiently repaired, leading to extensive bacterial death— even at sublethal concentrations of FQs (Figure 4, bottom right). HR is a conserved and essential repair mechanism in many bacterial species. Since FQs are broad-spectrum antibiotics effective against a wide range of Gram-positive and Gram-negative bacteria, the sensitizing effect of HR disruption may extend beyond mycobacteria. For instance, previous transposon sequencing studies in *Escherichia coli* have shown that exposure to low concentrations of ciprofloxacin significantly reduced frequencies of transposon insertion mutants containing insertion within several HR-related genes, such as *recA, recB*, and *recC* ^22^. Similarly, deletion of *recA* has been shown to increase quinolone sensitivity in other bacterial species, including *Pseudomonas aeruginosa* and *Staphylococcus aureus* ^23^. These observations reinforce the concept that targeting HR or HR defective strains can potentiate the bactericidal activity of FQs across diverse pathogens.

**Figure 4:**
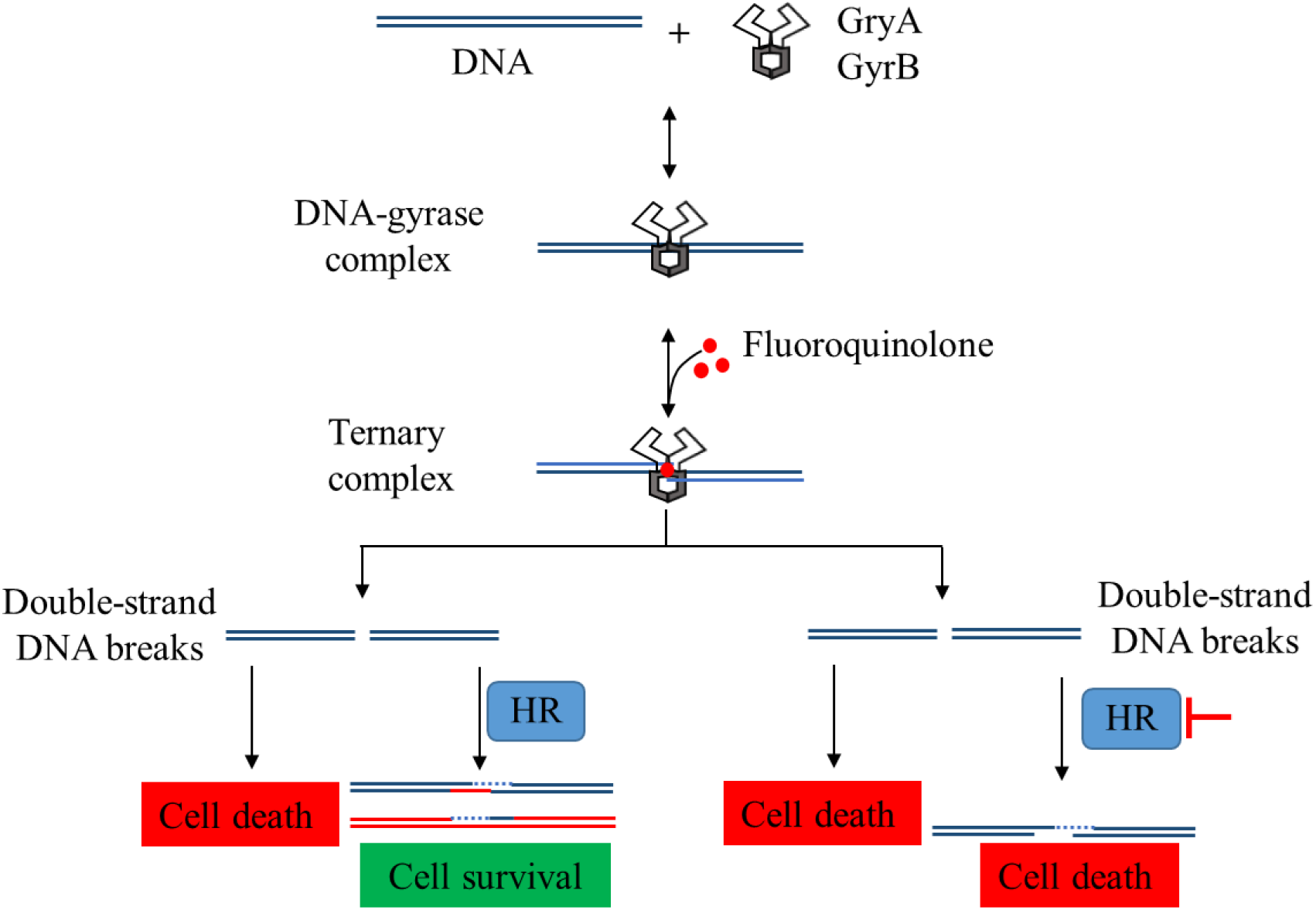
Scheme describing synergistic FQs sensitization by targeting the HR repair pathway. In wild-type mycobacteria, a portion of FQ-induced DSBs can be repaired by HR, allowing some cells to survive. However, when HR is disrupted, these DSBs remain unrepaired, rendering mycobacteria more sensitive to FQs.

Additionally, our *in vivo* data demonstrate that disruption of the HR repair pathway significantly impairs the survival of mycobacteria, consistent with prior studies indicating that DNA repair deficiencies impair mycobacterial proliferation under macrophage-simulated stress ^24^. These findings suggest that targeting key components of the HR pathway could both augment susceptibility mycobacteria to FQs and independently undermine their persistence within hosts. This dual mechanism underscores the therapeutic potential of HR inhibition and provides a compelling strategy to enhance the efficacy of existing antibiotics, offering new avenues for the treatment of multidrug-resistant mycobacterial infections.

Importantly, our study demonstrates that sensitizing mycobacteria to FQs is specific to HR disruption, deletion of genes involved in alternative repair pathways NHEJ and SSA did not affect FQ susceptibility. This specificity can be attributed to the mechanistic and fidelity differences among these DNA repair pathways. The HR pathway serves as a high-fidelity repair mechanism, crucial for accurately repairing DSBs during DNA replication by using a homologous sequence as a template, thereby maintaining genomic stability ^14^. In contrast, both SSA and NHEJ are more error-prone and often result in deletions or insertions that compromise genomic integrity ^14^. Therefore, disrupting SSA and NHEJ does not sensitize *M. abscessus* to FQs, rather, the mutations introduced by these repair pathways may hinder the bacteria’s survival. Furthermore, NHEJ is poorly functional in *M. abscessus, M. tuberculosis*, and *M. smegmatis*, as DSBs induced by CRISPR-Cas systems cannot be efficiently repaired without the introduction of *nrgA* from *M. marinum* ^25, 26^. This underscores the limited functionality of the NHEJ pathway in these mycobacteria. Consequently, disrupting NHEJ does not increase FQs sensitivity, further emphasizing the critical role of HR in determining the bacterium’s response to DSBs induced by FQ treatment.

There are significant differences between prokaryotic and eukaryotic HR repair mechanisms for DSBs. Prokaryotes primarily rely on the RecA protein, which facilitates homologous pairing and recombination of DNA. This repair process is relatively simple, involving fewer repair factors. In contrast, eukaryotes utilize a more complex repair network, involving multiple repair factors such as RAD51, BRCA1, and BRCA2, which coordinate the repair process through a tightly regulated network of interactions ^27^. Although functionally analogous proteins exist in both prokaryotes and eukaryotes, significant structural and biochemical differences have been observed. For example, the RecA ortholog in eukaryotes, Rad51, and its paralogs are functionally similar to RecA but differ structurally and biochemically, despite having a moderately similar ATP-binding domain ^28, 29^. Consequently, these distinctions suggest that inhibitors targeting RecA-mediated DNA binding should be specific to bacterial species. Thus, the divergence in HR pathways underscores their potential as targets to enhance the antibacterial efficacy of FQs.

Several compounds have demonstrated the ability to inhibit key enzymes in the HR repair pathway, particularly targeting RecA. For example, phthalocyanine tetrasulfonic acid has been shown to inhibit RecA activity, thereby enhancing the antibacterial efficacy of quinolones against both Gram-negative and Gram-positive bacteria. Furthermore, it has been shown to significantly increase the efficacy of FQs against *E. coli* infections in mouse models ^30^. RecA is regulated by intein, which can only be activated into functional proteins through the process of protein splicing from their initially inactive precursor proteins. Notably, the FDA-approved chemotherapeutic agent cisplatin has been shown to inhibit intein splicing in *M. tuberculosis*, thereby impairing RecA maturation and function ^31^. These findings pave the way for rational development of RecA-targeted adjuvants that could be co-administered with FQs to treat drug-resistant mycobacterial infections.

In conclusion, this study identifies HR repair as a key determinant of FQs efficacy and bacterial survival in *M. abscessus* and other mycobacterial pathogens. By elucidating the genetic basis of enhanced FQ susceptibility upon HR disruption, these findings lay the groundwork for novel therapeutic strategies targeting bacterial DNA repair pathways. The development of HR inhibitors represents a promising avenue to re-sensitize multidrug-resistant mycobacteria to FQ therapy. This work represents a significant step toward addressing the urgent challenge of drug-resistant mycobacterial infections and advancing more effective treatment options for patients afflicted by these notoriously resilient pathogens.

## Materials and Methods

### Bacterial strains, growth conditions, and antimicrobial agents

*E. coli* DH5α was cultured at 37°Cin Luria-Bertani (LB) broth and on LB agar plates. *M. abscessus* subsp. *abscessus* GZ002 ^32^, *M. tuberculosis* H37Ra, and their derivative strains were grown at 37°Cin Middlebrook 7H9 broth (Difco) supplemented with 10% oleic acid-albumin-dextrose-catalase (OADC, Difco) and 0.05% Tween 80, or on Middlebrook 7H10 agar (Difco) containing 10% OADC. Kanamycin, and zeocin were used as needed at final concentrations of 100 μg/mL, and 30 μg/mL, respectively.

### Generation of FQs-hypersensitive Tn mutants

Transposon mutant libraries of *M. abscessus* were constructed using *Himar1* mutagenesis, as previously described ^32^. Briefly, 100 mL of mid-log-phase *M. abscessus* culture, with an optical density at 600 nm (OD_600_) of approximately 0.7 to 1.0, was infected with 1 ∼ 2 × 10^11^ PFU/mL of MycoMarT7 phage at 37°Cfor 4 hours. Following infection, cells were pelleted, washed, and plated on Middlebrook 7H10 agar containing kanamycin. Tn mutants were then replica streaked onto Middlebrook 7H10 agar using a high-throughput bacteria/fungi screening workstation (Fluent 480, TECAN) with or without a sub-inhibitory concentration of LEV (2 μg/mL). Mutants showing growth impairment on LEV-containing plates were collected and subjected to MIC determination. The Tn insertion sites in the LEV-hypersensitive mutants were identified using whole-genome sequencing and confirmed by PCR ^32^.

### Gene knockout and complementation

Gene knockout in *M. abscessus* was performed using recombineering or CRISPR-assisted recombineering, following established protocols ^32^. For recombineering, an allelic exchange substrate for gene deletion was constructed by amplifying the upstream and downstream flanking arms and cloning them adjacent to a zeocin resistance gene, which was inserted into the pBluescript II SK (+) vector. These constructs were amplified by PCR and electroporated into freshly prepared electrocompetent *M. abscessus* cells carrying the recombineering plasmid pJV53. For CRISPR-assisted recombineering, the substrates, together with the crRNA-expressing pCRZeo plasmid, were electroporated into *M. abscessus* cells carrying the recombineering plasmid pJV53-Cpf1. Plates were incubated at 30°Cfor 5 days. Colonies were selected on zeocin-containing plates and verified by PCR and sequencing. Subsequently, complemented strains were constructed by reintroducing targeted genes and their homolog from *M. tuberculosis*. All primers used in this study are listed in Table S3.

### Antibiotics susceptibility testing

MICs were determined using the broth microdilution method. Briefly, mycobacterial cells were inoculated at a density of 5 × 10^5^ CFU/mL into 7H9 medium (without tween 80) containing 2-fold serial dilutions of the drugs. *M. abscessus* strains were incubated at 37°Cfor 14 days for clarithromycin and for 3 days for other drugs. The MIC was defined as the lowest antibiotic concentration that prevented visible bacterial growth. Herein, wild-type, knockout, and complement strains were propagated in 7H9 medium at 37°C until reaching an optical density (OD_600_ nm) of 0.6. Subsequently, ten-fold serial dilutions were spotted onto plain Middlebrook 7H10 agar (Drug-free serving as a control) alongside plates supplemented with varying FQs concentrations. Following a 3-day incubation at 37°C, the plates were examined. The experiment was performed in triplicate and repeated independently three times.

### Mouse infection and treatment

Following previously established protocols ^22^, female BALB/c mice (GemPharmatech Co., Ltd, - 8 weeks old) were infected with *M. abscessus* using a whole-body inhalation exposure system. To ensure appropriate immunosuppression, the mice were treated with dexamethasone (Sigma-Aldrich). Dexamethasone was dissolved in sterile 1 × phosphate-buffered saline (PBS, pH 7.4) and administered via daily subcutaneous injection at a dosage of 4 mg/kg/day, starting 1 week prior to infection and continuing throughout the experiment. Mice were sacrificed 4 hours post-infection to assess the initial bacterial burden in the lungs.

Subsequently, the mice received MFX at a dose of 100 mg/kg via oral gavage once daily, with PBS as the solvent. After 10 days of drug treatment, the mice were euthanized, and their lungs and spleens were aseptically removed, homogenized in PBS, and plated on Middlebrook 7H11 agar supplemented with 10% OADC. Colonies were counted after 3 days of incubation at 37°C. All animal care and experimental protocols were approved by the Institutional Animal Ethics Committee of the Guangzhou Institutes of Biomedicine and Health, Chinese Academy of Sciences.

## ACKNOWLEDGMENTS

This work was supported by the National Key R&D Program of China (2021YFA1300904), Postdoctoral Fellowship Program of CPSF (GZC20232688), Guangdong Provincial Basic and Applied Basic Research Fund (2024A1515012412), Major Project of Guangzhou National Laboratory (GZNL2024A01009), Guangzhou Science and Technology Plan-Youth (2024A04J4273), Doctoral ‘Sail’ Project, Guangzhou Institute of Respiratory Diseases, First Affiliated Hospital of Guangzhou Medical University (SKLRD-Z-202412, SKLRD-Z-202301, SKLRD-OP-202324), and Key Research and Development Program of Guangzhou (2025B01J3019).

## AUTHOR CONTRIBUTIONS

**Shuai Wang**: Conceptualization (equal); investigation (equal); methodology (equal); validation (equal); visualization (equal); formal analysis (equal); writing–original draft (equal); writing–review and editing (equal). **Md Shah Alam:** Conceptualization (equal); investigation (equal); methodology (equal); validation (equal); visualization (equal); formal analysis (equal); writing–original draft (equal); writing–review and editing (equal). **Buhari Yusuf**: Original draft (supporting) and writing–review (supporting). **Jingran Zhang**: Data curation (equal) and formal analysis (supporting). **Ziwen Lu**: Formal analysis (supporting); software (supporting). **Xiaofan Zhang**: Formal analysis (supporting); software (supporting). **H.M. Adnan Hameed**: Original draft (supporting) and writing–review (supporting). **Mst. Sumaia Khatun**: Formal analysis (supporting); validation (supporting). **Chunyu Li**: Formal analysis (supporting); validation (supporting). **Lijie Li**: Formal analysis (supporting); validation (supporting). **Aleksey A. Vatlin**: writing–review, and editing (supporting); validation (supporting). **Aweke Mulu Belachew**: Writing–original draft (supporting) and software (supporting). **Xirong Tian**: Formal analysis (supporting); and Resources (supporting), **Yamin Gao**: Resources (supporting), writing–review, and editing (supporting). **Cuiting Fang**: Validation (supporting), and writing–review (supporting). **Rogers Peng Zhang**: Formal analysis (supporting); validation (supporting). **Jun Li**: Writing–review and editing (supporting). **Xinyue Wang**: Formal analysis (supporting); validation (supporting). **Liqiang Feng**: Writing–review and editing (supporting). **Li Wan**: Writing–review and editing (supporting). **Tianyu Zhang**: Conceptualization (equal); project administration (lead); funding acquisition (lead); resources (supporting), validation (equal), and writing–review and editing (equal).

## ETHICS STATEMENT

Animal procedures were approved by the institutional animal care and use committee of the Guangzhou Institutes of Biomedicine and Health (number 2024003).

