## Supplementary file for "Targeting homologous recombination repair to potentiate fluoroquinolone efficacy in *Mycobacterium abscessus*"

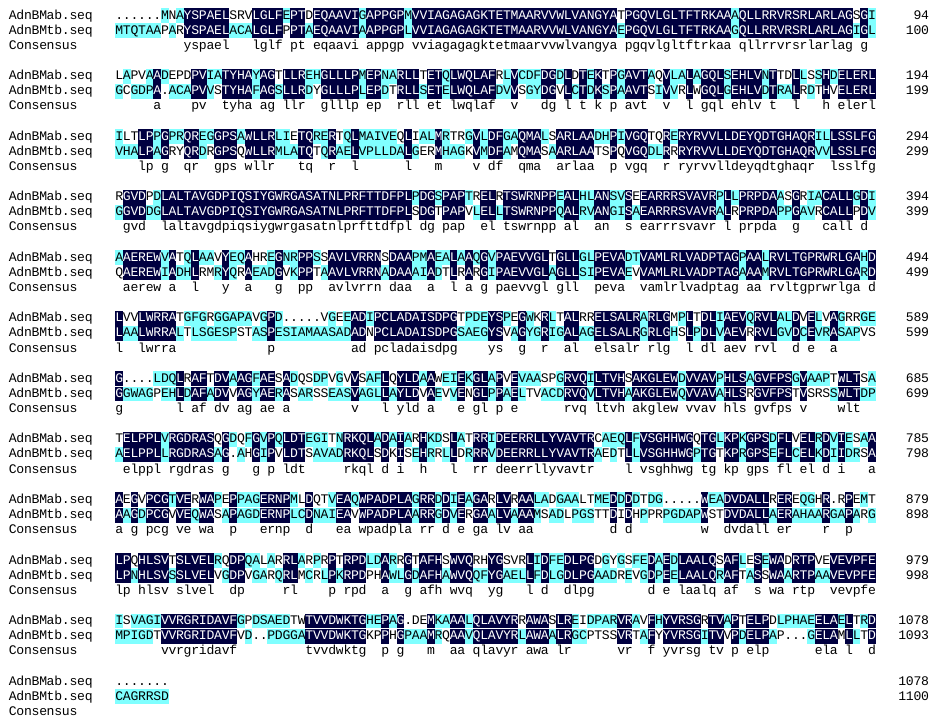

**Figure S1:** The alignment of amino acid sequences of AdnB proteins from *M. abscessus* (MAB_3515) and *M. tuberculosis*. *M. abscessus* showed 59.21% identity with AdnB of *M. tuberculosis*.

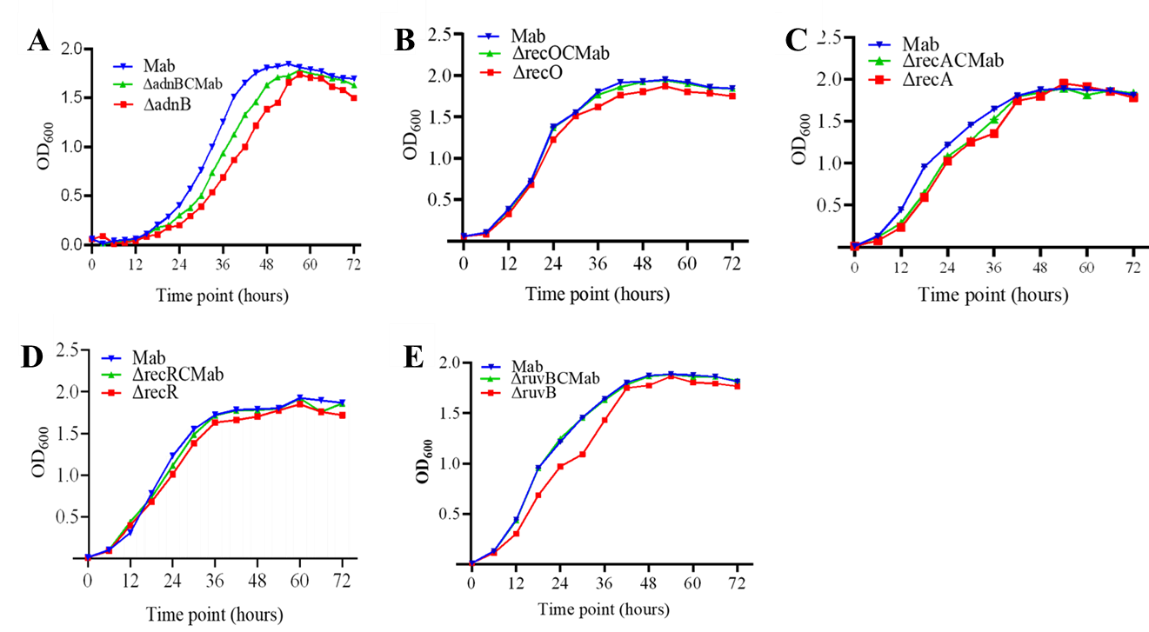

**Figure S2:** Growth curves of different *M. abscessus* strains. Growth curves were measured by OD₆₀₀ for the deletion mutants of (A) *adnB*, (B) *recO*, (C) *recA*, (D) *recR*, and (E) *ruvB* and the corresponding complementary strains with the wild *M. abscessus* (Mab) as a control under standard culture conditions.

**Table S1** MICs of various antibiotics to different *M. abscessus* strains

| Antibiotics^a^ | Strains/MIC (μg/mL) | | |
| --- | --- | --- | --- |
|  | Mab | S27 | ∆adnB |
| rifampicin | 128 | 128 | 128 |
| rifabutin | 8 | 8 | 8 |
| imipenem | 16-32 | 16-32 | 16-32 |
| vancomycin | 64-128 | 64-128 | 64-128 |
| clarithromycin | 64 | 64 | 64 |
| tigecycline | 2 | 2 | 2 |
| amikacin | 8 | 8 | 8 |
| linezolid | 64 | 64 | 64 |
| clofazimine | 2-4 | 2-4 | 2-4 |

^a^ The clarithromycin susceptibility results were recorded at 14 days due to its inducible resistance characteristics, while the results for other antibiotics were recorded at 3 days.

**Table S2** MICs of LEV and MXF for different *M. abscessus* strains

| Antibiotics | *M. abscessus* strains/ MICs (μg/mL) | | | |
| --- | --- | --- | --- | --- |
|  | Mab | Δku | ΔligD | ΔrecC |
| LEV | 16 | 16 | 16 | 16 |
| MFX | 8 | 8 | 8 | 8 |

**Table S3** Primers used in this study

| **Primers** | **Nucleotide sequences (5’ – 3’)** | **Description** |
| --- | --- | --- |
| adnBUP-F | CGCTCTAGAACTAGTGGATCCAAAGCCCTGCCACCAAAGA | Construction of the pBluescript II SK(+)-UadnBD plasmid containing the upstream and downstream homologous arms of *M. abscessus adnB* |
| adnBUP-R | TAGCCGTCGCAAGCTTGGAAGGCCAGCTGCCATAA |  |
| adnBDown-F | TTCCAAGCTTGCGACGGCTACGGAAGTTT |  |
| adnBDown-R | CTCGAGGTCGACGGTATCGATAGCGGCGTGATCACAAGGA |  |
| HindIIIdifzeo-F ^a^ | CCCAAGCTTAACGCCGATAAGACACATTATGTCAGTTGAATTCCGACCCGCACGAC | PCR amplification of zeocin resistance cassette with *dif* sites for constructing homologous recombination substrates |
| HindIIIdifzeo-R ^a^ | CCCAAGCTTAACTGACATAATGTGTCTTATCGGCGTTTCAGTCCTGCTCCTCGGCCA |  |
| adnBUZD-F | ATGGGGTGATATCGCCATAG | Amplification of homologous recombination substrates for construction of the *adnB* knockout mutant in *M. abscessus* |
| adnBUZD-R | AAAGCCCTGCCACCAAAGA |  |
| czadnB-F | GGCCAAGACAATTGCGGATCCGTCAGTCGTGAATGCATAC | Construction of a complementation plasmid for ∆adnBCMab |
| czadnB-R | GTTAACTACGTCGACATCGATACCAGGACAATCAGGAACA |  |
| recOUP-F | CTCGAGGTCGACGGTATCGATCAGCTCGCAGGTGTTGGTCT | Construction of the pBluescript II SK(+)-UrecOD plasmid containing the upstream and downstream homologous arms of *M. abscessus recO* |
| recOUP-R | TAGCTTGCGTTGACGGGTCAGCAGGGTCACG |  |
| recODown-F | TGACCCGTCAACGCAAGCTACGGACACTC |  |
| recODown-R | CGCTCTAGAACTAGTGGATCCGTAGACGCTGCTCCACAACC |  |
| recAUP-F | CGCTCTAGAACTAGTGGATCCTGGTGGGGGACACGGCTTT | Construction of the pBluescript II SK(+)-UrecAD plasmid containing the upstream and downstream homologous arms of *M. abscessus recA* |
| recAUP-R | CATATCGCCGGTCGGGATCACGGAGAT |  |
| recADown-F | TGATCCCGACCGGCGATATGGGCGTGGAGCAG |  |
| recADown-R | CTCGAGGTCGACGGTATCGATCCGAGGCGAGGGCATTCTT |  |
| ruvBUP-F | CGCTCTAGAACTAGTGGATCCGCAGAGGAACAACGGAAGG | Construction of the pBluescript II SK(+)-UruvBD plasmid containing the upstream and downstream homologous arms of *M. abscessus ruvB* |
| ruvBUP-R | AACTGCGTGTCAGGGCGGGTGGGCCGGATAACAGG |  |
| ruvBDown-F | ACCCGCCCTGACACGCAGTTTCG |  |
| ruvBDown-R | CTCGAGGTCGACGGTATCGATTTCGGGGGATGTGAGCAAA |  |
| recRUP-F | CTCGAGGTCGACGGTATCGATACCGTCACCGATGCCGCTGTT | Construction of the pBluescript II SK(+)-UrecRD plasmid containing the upstream and downstream homologous arms of *M. abscessus recR* |
| recRUP-R | ATCCGCACCAAGTAGTCCTGGACCGGGCCCTCAAA |  |
| recRDown-F | CAGGACTACTTGGTGCGGATGCTGC |  |
| recRDown-R | CGCTCTAGAACTAGTGGATCCTGATCGACTTCGAGGTGGAA |  |
| recCUP-F | CGCTCTAGAACTAGTGGATCCACGCCCTGAGCGCCGCAGTT | Construction of the pBluescript II SK(+)-UrecCD plasmid containing the upstream and downstream homologous arms of *M. abscessus recC* |
| recCUP-R | TCGTTTCCCCCTCGGTCTCTCGCACGCCCGCACACACCC |  |
| recCDown-F | AGAGACCGAGGGGGAAACGAA |  |
| recCDown-R | CTCGAGGTCGACGGTATCGATAGGCGCAATGCCGCCGGATAG |  |
| kuUP-F | CGCTCTAGAACTAGTGGATCCAGGCCCTTGCTTCCACTGG | Construction of the pBluescript II SK(+)-UkuD plasmid containing the upstream and downstream homologous arms of *M. abscessus ku* |
| kuUP-R | ATAGGTGTCGGTGCGTTTGTAGCGAATCCG |  |
| kuDown-F | ACAAACGCACCGACACCTATCAGTCACAGCTGC |  |
| kuDown-R | CTCGAGGTCGACGGTATCGATCCACTACTTCAGCCAAATCCGT |  |
| ligDUP-F | CTCGAGGTCGACGGTATCGATCCAGCAGCACATACGATTTG | Construction of the pBluescript II SK(+)-UligDD plasmid containing the upstream and downstream homologous arms of *M. abscessus ligD* |
| ligDUP-R | CGGTCCTTCTCGAAGAACGAC |  |
| ligDDown-F | TCGTTCTTCGAGAAGGACCGCCGCAGCAGCGGAATAGGG |  |
| ligDDown-R | CGCTCTAGAACTAGTGGATCCTTCGTCGCGTGAGCGAGCAC |  |
| recO-crRNA-F | AT CCCTGCACCGCCTCACCGTTGGTGA | Construction of the pCRZeo-recO plasmid containing a crRNA targeting *M. abscessus recO* |
| recO-crRNA-R | AGCTTCACCAACGGTGAGGCGGTGCAGGGATCT |  |
| recA-crRNA-F | ATGGGTGAACCGAACATCACGCA | Construction of the pCRZeo-recA plasmid containing a crRNA targeting *M. abscessus recA* |
| recA-crRNA-R | AGCTTGCGTGATGTTCGGTTCACCCATCT |  |
| recR-crRNA-F | ATCATTTGTTGGGTGTCGAGGCGCA | Construction of the pCRZeo-recR plasmid containing a crRNA targeting *M. abscessus recR* |
| recR-crRNA-R | AGCTTGCGCCTCGACACCCAACAAATGATCT |  |
| ruvB-crRNA-F | ATGAGCTCGTCGGGCTCGTAGAA | Construction of the pCRZeo-ruvB plasmid containing a crRNA targeting *M. abscessus ruvB* |
| ruvB-crRNA-R | AGCTTTCTACGAGCCCGACGAGCTCATCT |  |
| recC-crRNA-F | ATCGCGCCCTGGACTACACCCTGCA | Construction of the pCRZeo-recC plasmid containing a crRNA targeting *M. abscessus* *recC* |
| recC-crRNA-R | AGCTTGCAGGGTGTAGTCCAGGGCGCGATCT |  |
| ku-crRNA-F | ATCTGGTCCAACACCGGGAAGTCGA | Construction of the pCRZeo-ku plasmid containing a crRNA targeting *M. abscessus* *ku* |
| ku-crRNA-R | AGCTTCGACTTCCCGGTGTTGGACCAGATCT |  |
| ligD-crRNA-F | ATGTGGATTGGAGCCAGAACAGCA | Construction of the pCRZeo-ligD plasmid containing a crRNA targeting *M. abscessus ligD* |
| ligD-crRNA-R | AGCTTGCTGTTCTGGCTCCAATCCACATCT |  |
| czadnB-F | GGCCAAGACAATTGCGGATCCGTCAGTCGTGAATGCATAC | Construction of the complementation plasmid carrying the *M. abscessus adnB* gene |
| czadnB-R | GTTAACTACGTCGACATCGATACCAGGACAATCAGGAACA |  |
| czadnB-Mtb-F | GGCCAAGACAATTGCGGATCCATGACCCAAACCGCGGCA | Construction of the complementation plasmid carrying the *M. tuberculosis adnB* gene |
| czadnB-Mtb-R | GTTAACTACGTCGACATCGATTCAGGTGTCCGACCGCCG |  |
| czrecO-F | GGCCAAGACAATTGCGGATCCATGCGGCTGTATCGGGACC | Construction of the complementation plasmid carrying the *M. abscessus recO* gene |
| czrecO-R | GTTAACTACGTCGACATCGATCTAGCCACCGGCTATATCCTGC |  |
| czrecO-Mtb-F | GTTAACTACGTCGACATCGATGCGAACCCGCTACAGAAAC | Construction of the complementation plasmid carrying the *M. tuberculosis recO* gene |
| czrecO-Mtb-R | GGCCAAGACAATTGCGGATCCCGACGAAACGGAGAACACC |  |
| czrecA-F | GGCCAAGACAATTGCGGATCCCGACGAAACGGAGAACACC | Construction of the complementation plasmid carrying the *M. abscessus recA* gene |
| czrecA-R | GTTAACTACGTCGACATCGATGCGAACCCGCTACAGAAAC |  |
| czrecA-Mtb-F | GGCCAAGACAATTGCGGATCCATGACGCAGACCCCCGAT | Construction of the complementation plasmid carrying the *M. tuberculosis recA* gene |
| czrecA-Mtb-R | GTTAACTACGTCGACATCGATTCAGAAGTCGACGGGGGC |  |
| czrecR-F | GGCCAAGACAATTGCGGATCCGTGTTTGAGGGCCCGGTCCA | Construction of the complementation plasmid carrying the *M. abscessus recR* gene |
| czrecR-R | GTTAACTACGTCGACATCGATCGGGTCGCCGCCCCCTCTGA |  |
| czrecR-Mtb-F | GGCCAAGACAATTGCGGATCCATGTTTGAGGGACCCGTCCA | Construction of the complementation plasmid carrying the *M. tuberculosis recR* gene |
| czrecR-Mtb-R | GTTAACTACGTCGACATCGATTCAGGCGAGCACGCGGCGG |  |
| czruvB-F | GGCCAAGACAATTGCGGATCCCGCAAAACAAGCCGAGCAA | Construction of the complementation plasmid carrying the *M. abscessus ruvB* gene |
| czruvB-R | GTTAACTACGTCGACATCGATGTACAGGCGAGGGCAAGCA |  |
| czruvB-Mtb-F | GGCCAAGACAATTGCGGATCCATGACCGAGCGGTCCGAC | Construction of the complementation plasmid carrying the *M. tuberculosis ruvB* gene |
| czruvB-Mtb-R | GTTAACTACGTCGACATCGATCTACTCGAACAACCCCGGTTG |  |

^a^ The *dif* core sites are underlined.
